## Supplementary figures and images for "Projection-specific integration of convergent thalamic and retrosplenial signals in the presubicular head direction cortex"

### Figure 1 - figure supplement 1

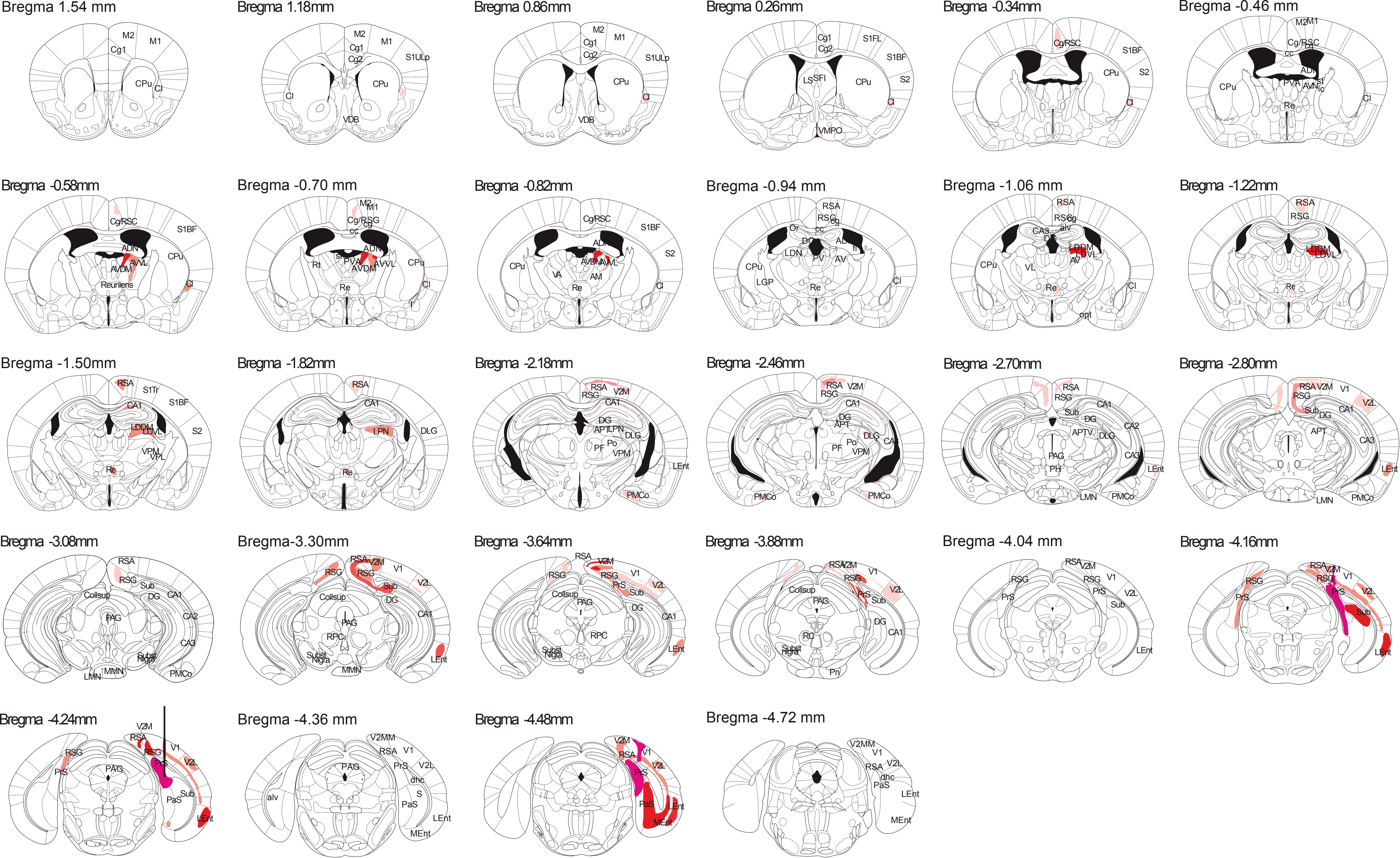

### Figure 1 - figure supplement 2

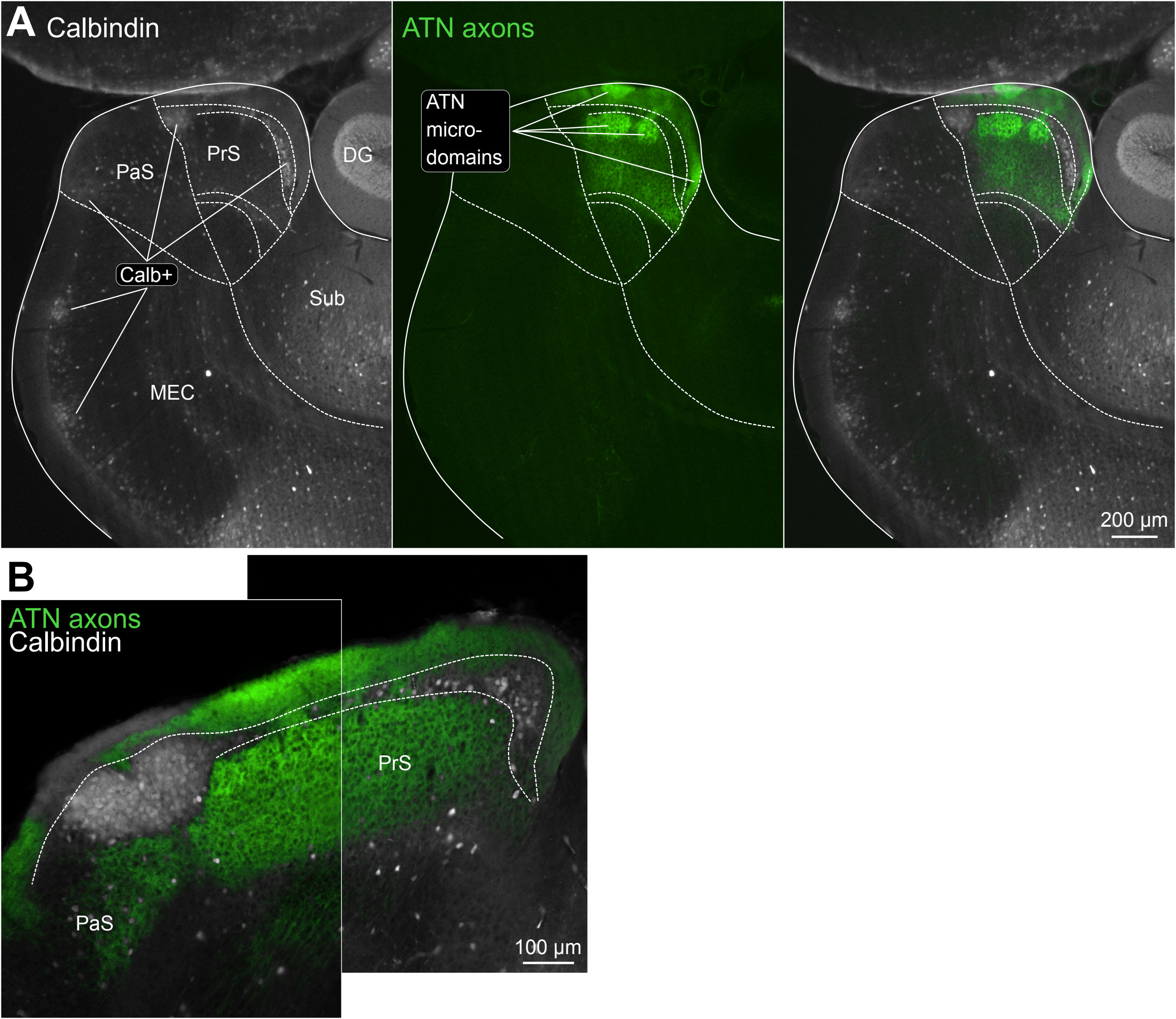

### Figure 2 - figure supplement 2

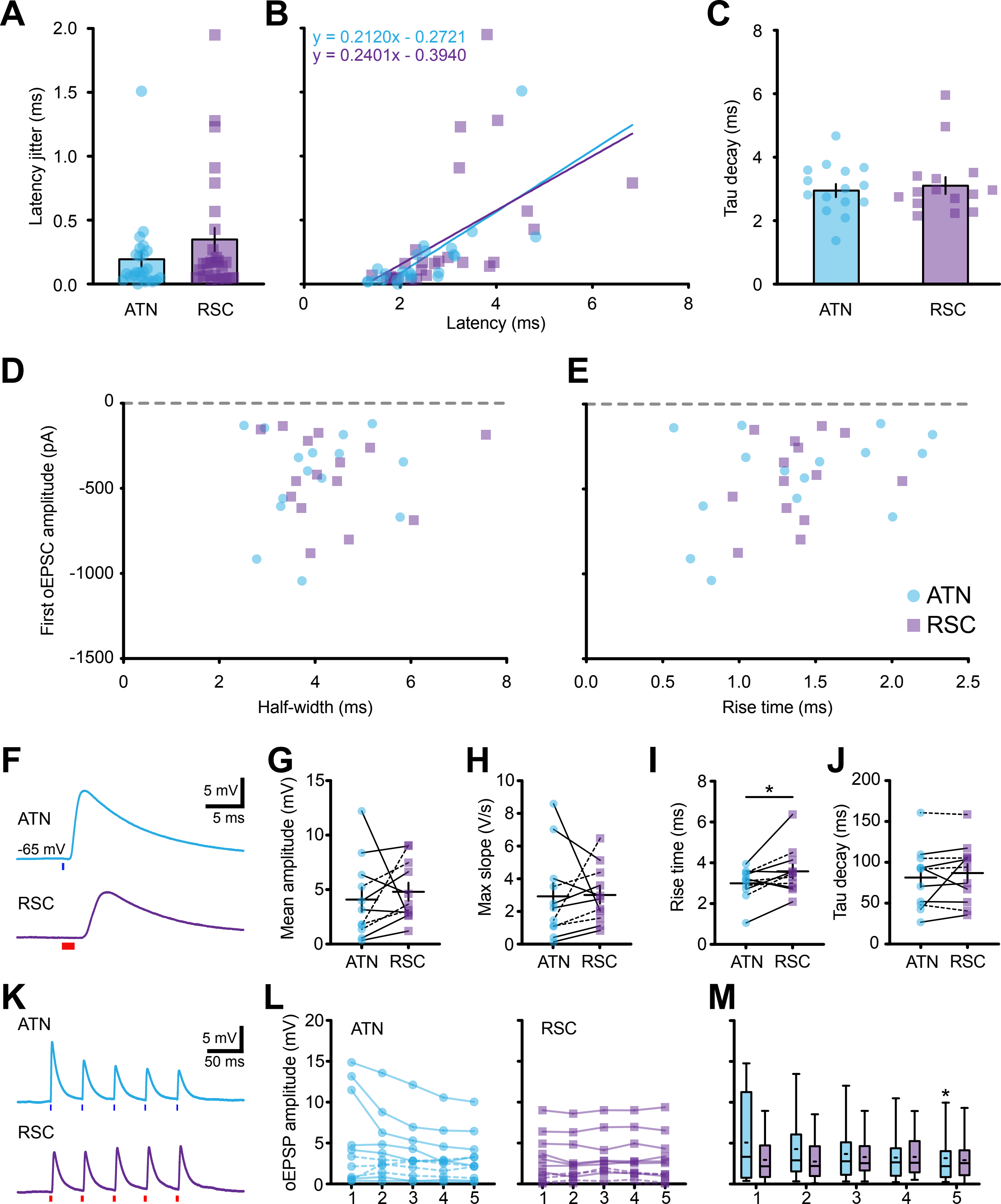

### Figure 6 - figure supplement 1

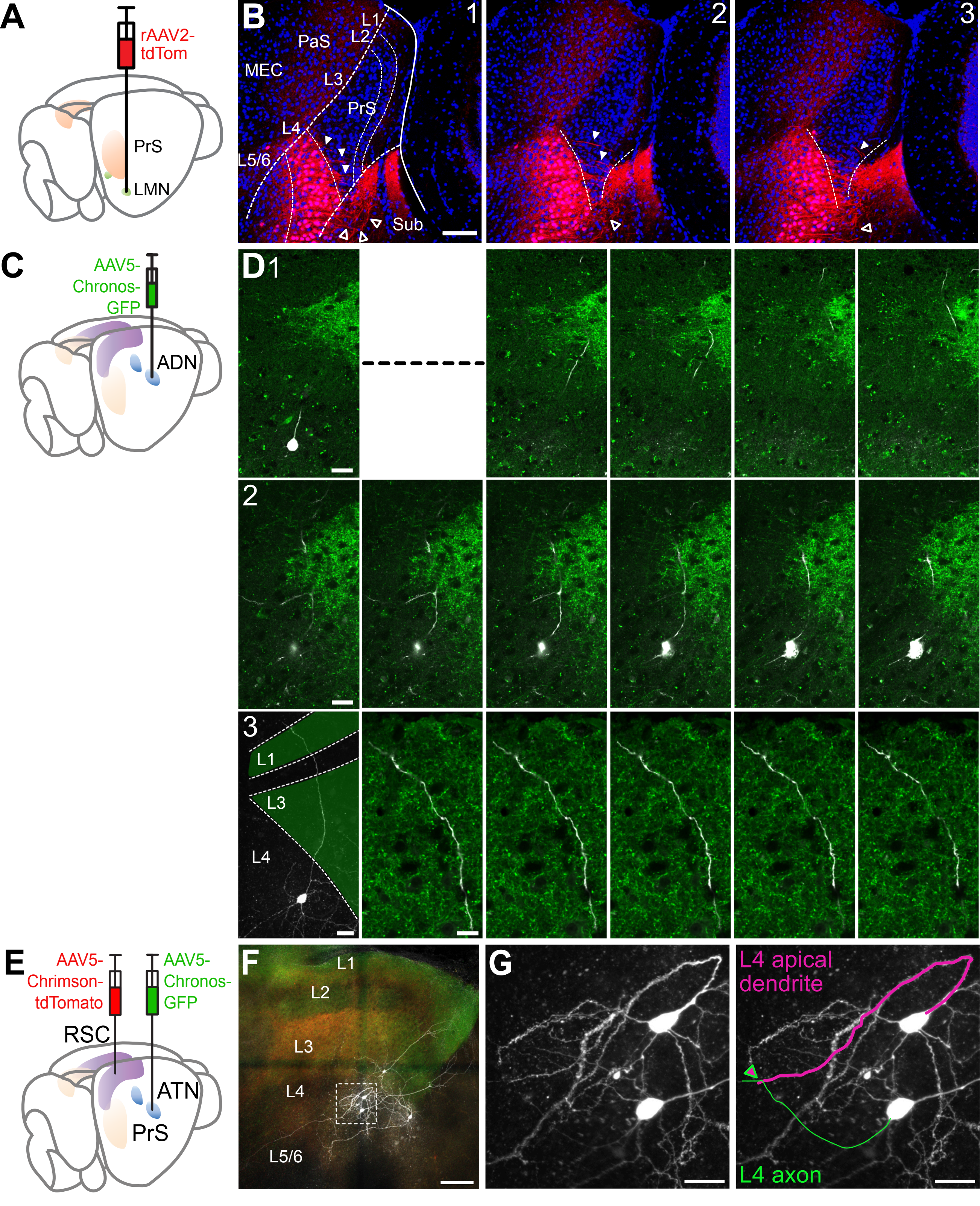

### Figure 6 - figure supplement 2

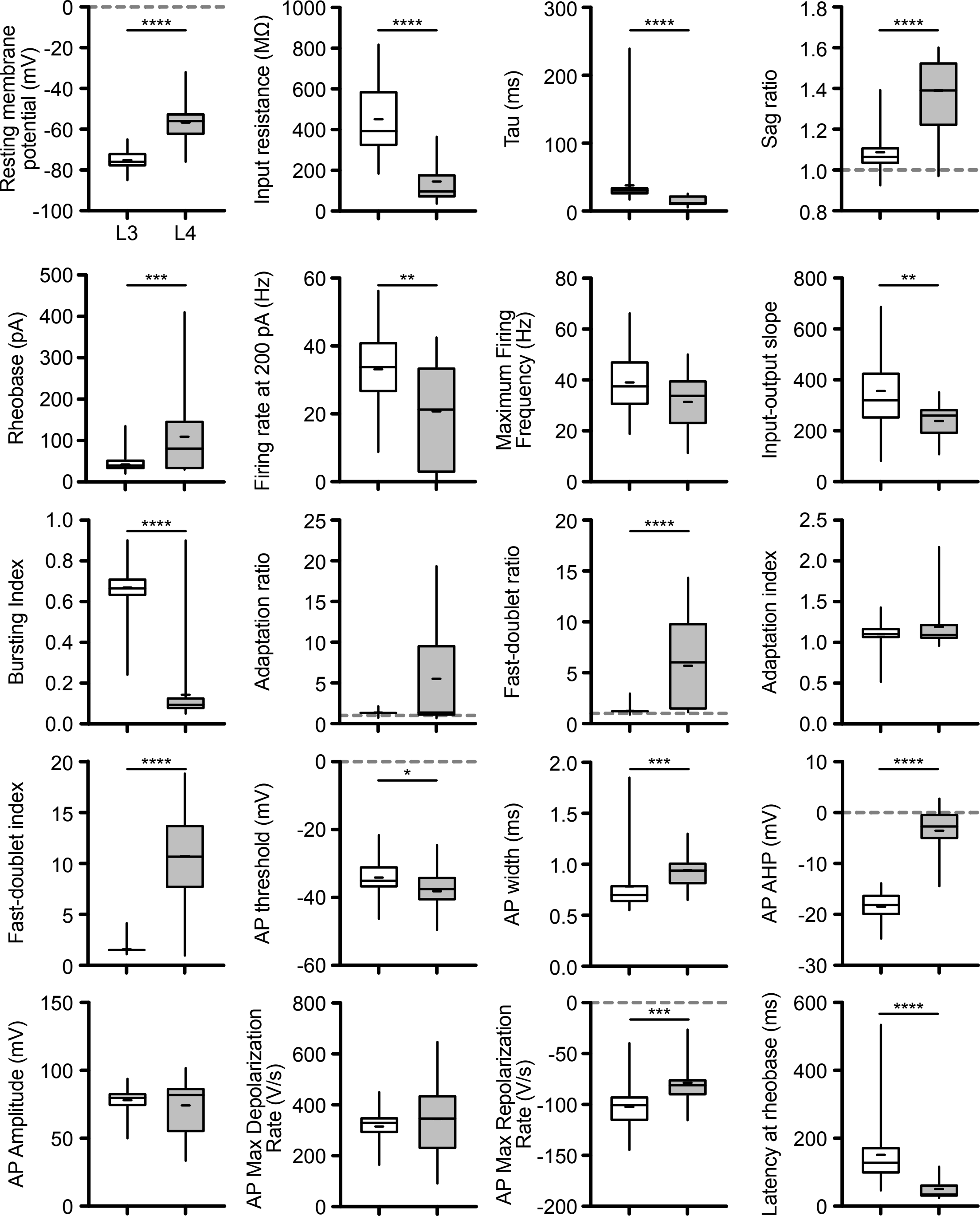

### Figure 7 - figure supplement 1

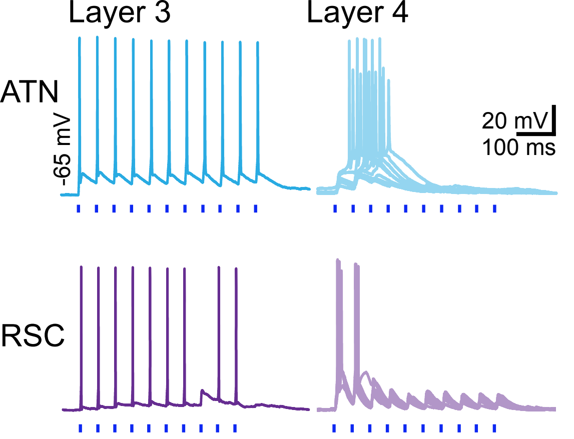

### Figure 7 - figure supplement 2

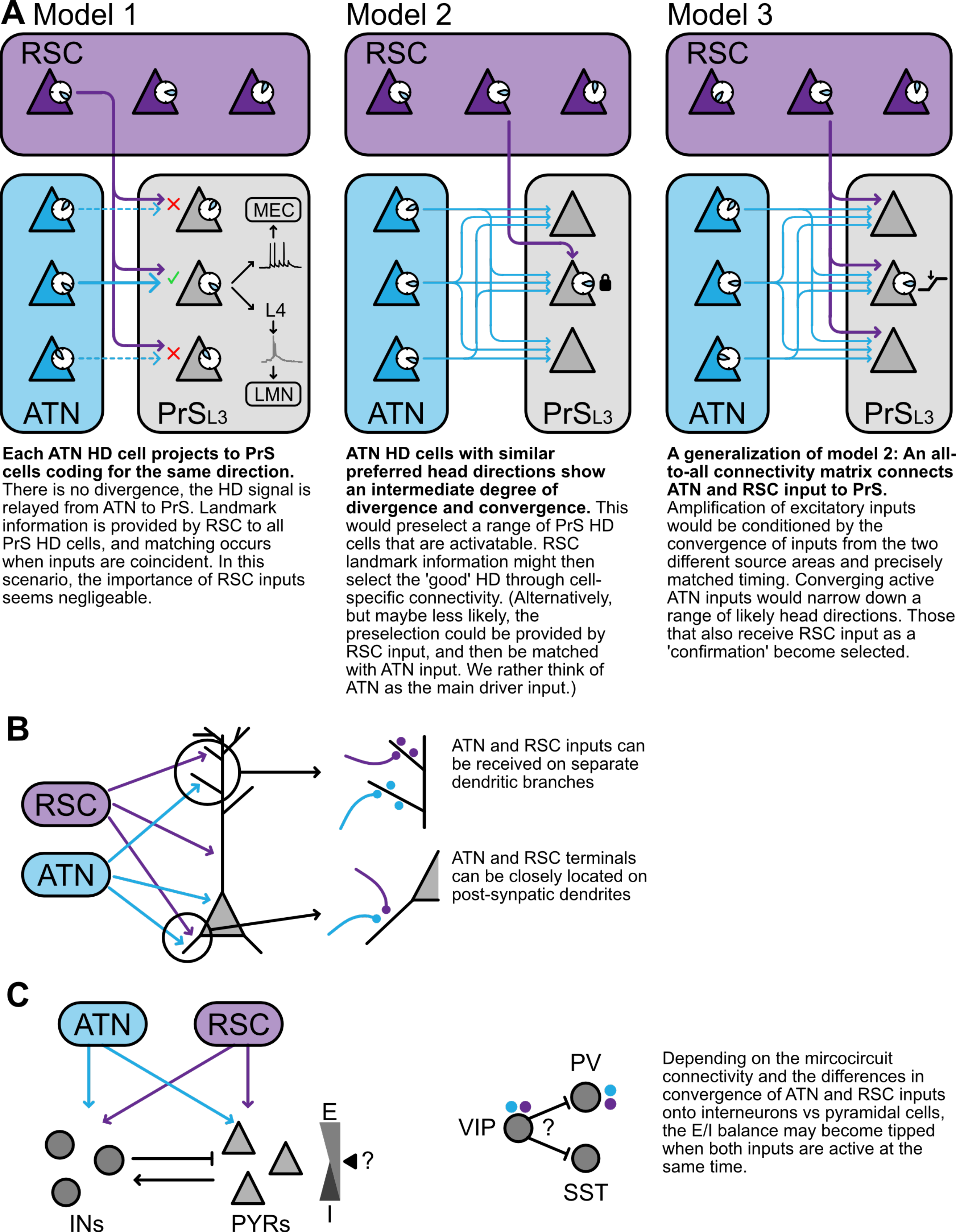
