## Supplementary material for "Projection-specific integration of convergent thalamic and retrosplenial signals in the presubicular head direction cortex": Figure 2 - figure supplement 1

| | ATN | | | RSC | | | *p-value* |
|---|---|---|---|---|---|---|---|
| | Mean | SEM | n | Mean | SEM | n | Mann-Whitney |
| **Resting membrane potential (mV)** | -71.37 | 1.85 | 27 | -73.50 | 1.52 | 38 | *ns* |
| **Neuronal input resistance (MΩ)** | 414.5 | 31.5 | 27 | 349.8 | 23.4 | 38 | *ns* |
| **Tau 1 (ms)** | 26.46 | 1.80 | 27 | 22.25 | 1.19 | 38 | *0.0481* |
| **Sag ratio at -100 mV** | 1.07 | 0.01 | 27 | 1.10 | 0.01 | 38 | *ns* |
| **Rheobase current (pA)** | 51.88 | 5.25 | 27 | 63.00 | 3.67 | 38 | *0.0188* |
| **Firing rate at 200 pA (Hz)** | 39.21 | 3.87 | 27 | 45.10 | 3.02 | 38 | *ns* |
| **Maximum firing frequency (Hz)** | 48.19 | 4.56 | 27 | 64.84 | 5.02 | 38 | *0.0035* |
| **Input-output slope (Hz/nA)** | 357.5 | 30.0 | 27 | 376.7 | 24.6 | 38 | *ns* |
| **AP threshold (mV)** | -34.13 | 0.86 | 27 | -33.48 | 0.91 | 38 | *ns* |
| **AP width (ms)** | 0.77 | 0.06 | 27 | 0.68 | 0.04 | 38 | *ns* |
| **AP AHP (mV)** | -20.25 | 0.64 | 27 | -18.77 | 0.47 | 38 | *ns* |
| **AP rise amplitude (mV)** | 77.84 | 1.67 | 27 | 79.77 | 1.26 | 38 | *ns* |
| **AP maximum depolarization rate (V/s)** | 323.7 | 11.0 | 27 | 350.7 | 9.2 | 38 | *0.0326* |
| **AP maximum repolarization rate (V/s)** | -107.9 | 6.1 | 27 | -121.7 | 5.9 | 38 | *ns* |
| **Onset latency at rheobase (ms)** | 181.2 | 24.3 | 27 | 139.4 | 17.6 | 38 | *ns* |
