## Supplementary material for "Projection-specific integration of convergent thalamic and retrosplenial signals in the presubicular head direction cortex": Figure 3 - figure supplement 1

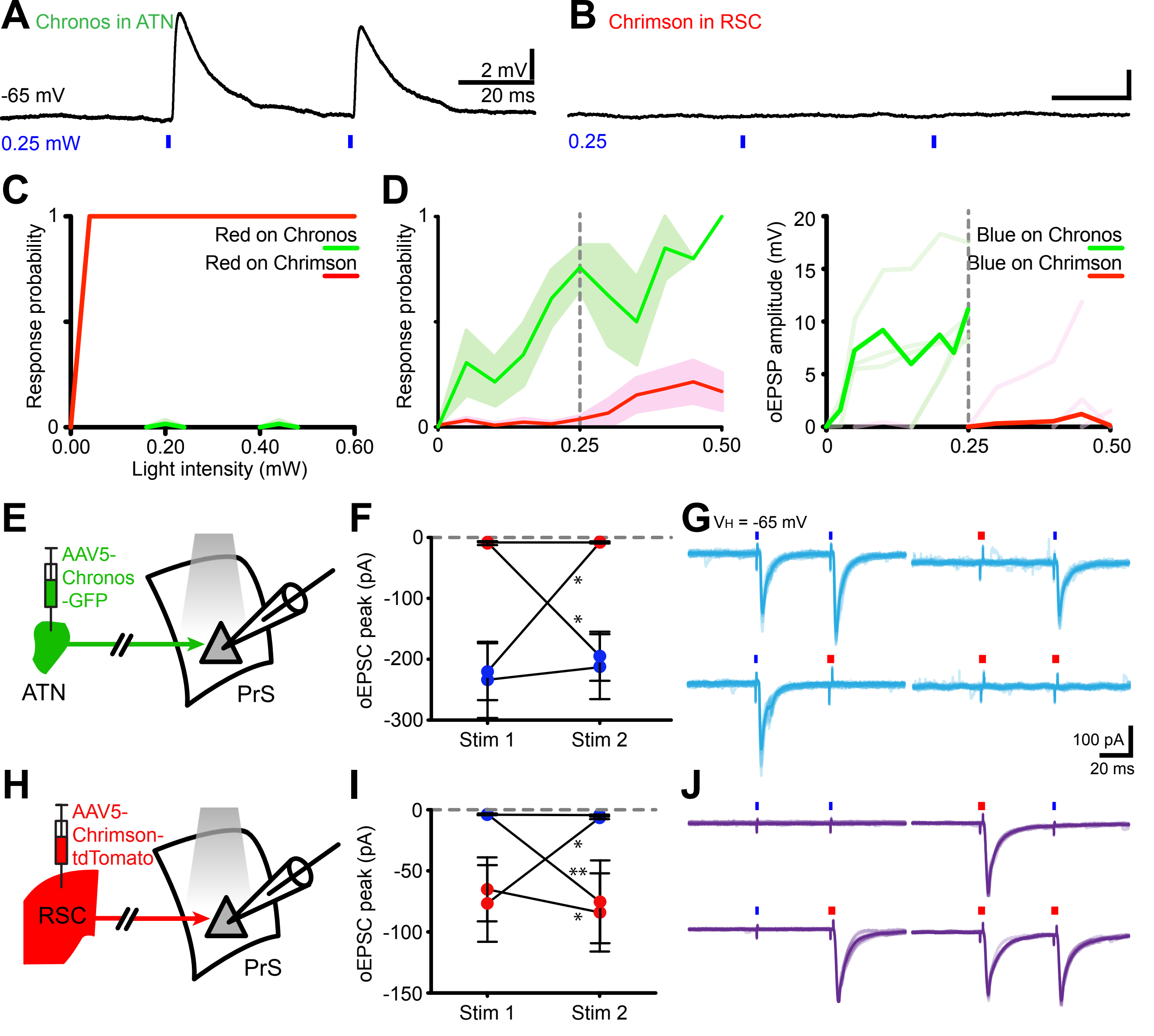


**K**

| | | Stim 1 Mean amplitude (pA) | Stim 1 SEM | Stim 2 Mean amplitude (pA) | Stim 2 SEM | n | *Wilcoxon test* |
|---|---|---|---|---|---|---|---|
| **Chronos in ATN** | Blue - Blue | -233.8 | 62.7 | -212.1 | 53.3 | 6 | *ns* |
| | Blue - Red | -220.1 | 46.7 | -7.0 | 0.7 | 6 | *0.0312* |
| | Red - Red | -9.5 | 3.2 | -8.2 | 1.7 | 6 | *ns* |
| | Red - Blue | -6.8 | 0.4 | -194.4 | 40.1 | 6 | *0.0312* |
| **Chrimson in RSC** | Blue - Blue | -4.1 | 0.3 | -4.4 | 0.5 | 8 | *ns* |
| | Blue - Red | -3.7 | 0.6 | -75.3 | 33.8 | 8 | *0.0156* |
| | Red - Red | -65.1 | 26.2 | -84.0 | 32.0 | 8 | *0.0234* |
| | Red - Blue | -76.7 | 31.4 | -6.6 | 1.0 | 8 | *0.0078* |
